# Progressive Remodeling of Global Protein Interaction Networks in a Mouse Model of Tauopathy

**DOI:** 10.1101/2025.07.05.663296

**Authors:** Weiwei Lin, Sadhna Phanse, Sophie J.F. van der Spek, Noah Lampl, Morgan C. Stephens, Alejandro Rondon Ortiz, Alexandria Taylor, Ryan Hekman, Rebecca Roberts, Lulu Jiang, Pierre Havugimana, Juan Botas, Andrew Emili, Benjamin Wolozin

## Abstract

Neurodegenerative disease is marked not just by loss of proteins or cells, but by dynamic rewiring of macromolecular interaction networks that precede and drive pathology. Here, we present the first temporally resolved, systems-scale map of multi-protein complex remodeling in a tauopathy model, integrating co-fractionation mass spectrometry, quantitative phosphoproteomics, and machine learning to decode phosphorylation-dependent shifts in protein interactomes across disease progression. This interactomic atlas identifies functionally validated assemblies—including MAPT-Dpysl2 and Cyfip1-actin complexes—that modulate early disease phenotypes in vivo. By revealing how phosphorylation tunes macromolecular complex architecture and function, this work reframes tauopathy as a disease of dynamic network instability, and establishes a generalizable framework for early detection and mechanistic dissection of neurodegeneration.

## INTRODUCTION

Alzheimer’s disease and related dementias (ADRD) are progressive neurodegenerative disorders characterized by synaptic failure, neuroinflammation, and eventual neuronal loss. While the accumulation of amyloid-β plaques and hyperphosphorylated microtubule-associated protein tau (MAPT) tangles define the pathological hallmark of ADRD, the resulting pathophysiology impacts a broad spectrum of biological processes. Omics-based studies have identified the widespread disruptions in the brain’s proteomic architecture, including perturbations in multi-protein complexes that act as proximate drivers of neuronal dysfunction ^1-6^. However, translating measurements of protein levels to the functional remodeling of protein complexes remains challenging as these assemblies often contain multiple subunits that can be modulated by both post-translational modifications and by interactions with RNA, metabolites, or lipids. Proximity labeling is a powerful method to study local interactions, but it requires pre-defined baits for effective application^7^. To overcome these limitations, we recently developed an algorithm, termed EPIC, which infers protein complexes from biochemical co-fractionation profiles, and applied it to identify soluble complexes in a comprehensive manner ^8^. This effort culminated in the creation of BrainMap, which is a systems-level platform that couples deep chromatographic fractionation with multiplex mass spectrometry (mCF-MS) and machine learning to resolve native protein complexes from brain tissue ^6^.

In the present study, we apply and optimize the BraInMap pipeline to the PS19 transgenic mouse model of tauopathy, which expresses mutant human P301S MAPT and exhibits age-dependent MAPT accumulation, hyperphosphorylation, and neurodegeneration ^9,10^. MAPT is a microtubule-associated protein critical for cytoskeletal stability, axonal transport, and synaptic plasticity ^11,12^. In tauopathies, MAPT becomes pathologically hyperphosphorylated and mislocalized, forming toxic neurofibrillary tangles. By integrating mCF-MS with quantitative proteomics and phosphoproteomics across a longitudinal time course, we systematically chart the reorganization of soluble brain protein complexes during pathological progression. This work provides a global view of how endogenous brain complexes evolve across disease stages. We identified and tracked changes in 4072 soluble protein complexes in P301S MAPT versus wild-type (WT) mice at 3, 6 and 9 months of age. Systematic analysis of two representative complexes demonstrates validation of predicted interactions by immunoprecipitation–mass spectrometry. We further assess their functional importance in well-established *Drosophila* models expressing either wild-type human MAPT or secreted Aβ ^13-16^. Notably, MAPT-expressing flies exhibit pronounced sensitivity to targeted disruption of complex components, underscoring their relevance to tau-induced phenotypes..

We also explored the evolution of additional complexes over the disease course. One complex, containing MAPT and the axonal maintenance factor Dpysl2, exhibits decreased association with actin-binding proteins (e.g. actg1) and increased association with stress response proteins (Eno1, Aco2 and Prdx1/2). In contrast, Cyfip, which couples actin with the RNA binding protein FMR, gains new associations with key actin regulators (Abi1/2, Wasf2) during disease progression. Integrated phosphoproteomic analysis reveals dynamically altered phosphorylation events at specific residues that correlate with disease stage, suggesting regulatory relevance.

Together, these findings establish a dynamic, high-resolution framework for proteome disorganization in tauopathy and nominate specific macromolecular assemblies as putative candidate effectors and therapeutic targets in ADRD. By integrating longitudinal co-fractionation mass spectrometry and phosphoproteomics, this study maps the evolving global architecture of macromolecular complexes and provides mechanistic insight into how MAPT disrupts cellular homeostasis at the systems level.

## RESULTS

### Progressive MAPT pathology and neuronal degeneration in the PS19 P301S MAPT mouse model

To characterize the spatiotemporal progression of tauopathy, we analyzed brain tissue from PS19 P301S MAPT transgenic mice (PS19), which express human P301S mutant MAPT and recapitulate key features of tau-driven neurodegeneration^1^. As summarized in Figure 1a, we applied a multi-modal profiling strategy at 3, 6, and 9 months of age-spanning early, intermediate, and advanced stages of pathology. The PS19 model is well-established for its age-dependent accumulation of hyperphosphorylated and aggregated tau, with neurofibrillary tangle formation and progressive neuronal loss most prominently affecting the hippocampus and later extending into cortical regions^1^. By 6 months, the PS19 brains show marked accumulation of MC1-positive misfolded MAPT in the cortex, which intensify further by 9 months (Suppl. Fig. 1a&b, *red*). There is a corresponding decline in MAP2 staining in 6- and 9-month PS19 brains consistent with dendritic degeneration and synaptic loss, hallmark features of tau-mediated neuronal dysfunction (Suppl. Fig. 1a&b, *green*). These results confirm that our PS19 mouse cohort exhibits progressive MAPT pathology and associated neuronal compromise over time, providing a robust foundation for subsequent interactomic and (phospho) proteomic analyses.

**Figure 1.**
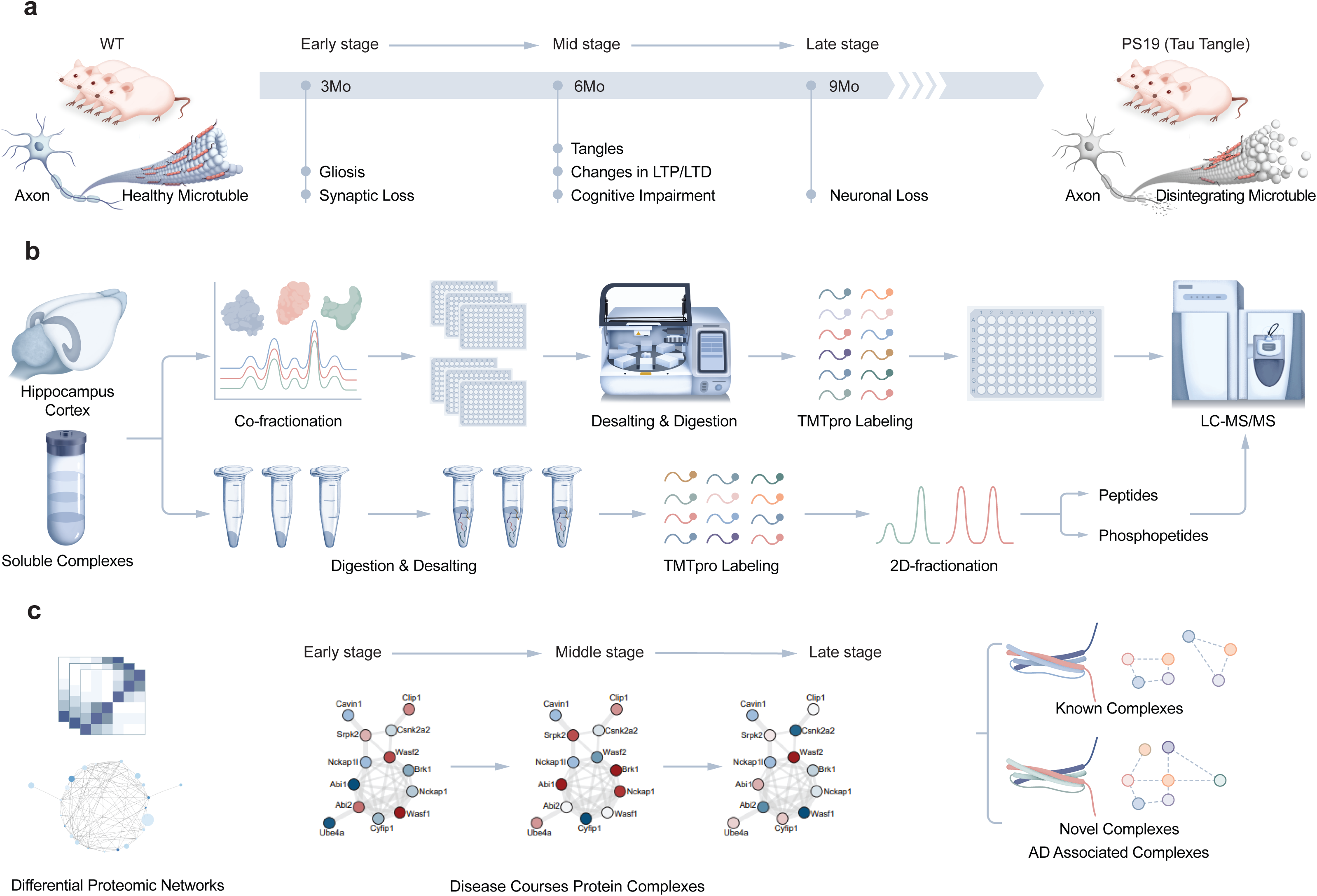
Schematic of multi-omic workflow to investigate AD-associated protein interactomes. (**a**) Brain tissues from 3-, 6-, and 9-month-old PS19 transgenic mice overexpressing human mutant tau, which present hallmark tauopathy features, were harvested. Hippocampus and cortex, regions vulnerable to MAPT aggregation, were isolated for protein extraction. (**b**) A gentle lysis protocol preserved native protein interactions. Identical lysates were used for global proteomics, phosphoproteomics, and mCF/MS. Peptides from each approach were labeled using Tandem Mass Tags (TMT) for quantification. (**c**) Differentially expressed proteins and phosphosites were integrated with machine-learning–predicted protein complexes (EPIC) to construct a global interactome associated with tauopathy.

### Proteomic profiling confirms early molecular signatures of tauopathy

To establish a molecular baseline and provide a comparative framework for our BraInMap (mCF-MS) interactome analysis, we first performed bulk proteomics on hippocampus and cortex, the two regions vulnerable to MAPT pathology, at early (3-month), mid-stage (6-month) and late-stage (9-month) timepoints in PS19 transgenic mice (**Fig. 1a & Suppl. Fig. 1c**). Tissue lysates were digested and labeled with isobaric tandem mass tags (TMT-16plex) coupled with offline fractionation to enhance depth of coverage prior to deep quantitative proteomic profiling via Orbitrap LC-MS (**Fig. 1b**).

Across all samples, we identified and quantified 8,397 proteins (**Supplementary Table 1**). As expected, human P301S MAPT was robustly detected across all PS19 timepoints (**Suppl. Fig. 1d**). In addition, at 3 months, we detected 180 significantly upregulated and 197 downregulated mouse proteins (p <0.05, FC >1.2; **Suppl. Fig. 2a**). We then compared our entire list of 8,379 identified proteins with known ADRD-linked human genes reported by GWAS^2^. This comparison revealed 719 overlapping proteins. Of these 719, a subset of 161 proteins showed significant changes in our 3-month PS19 brains compared with the wild-type (<0.05, FC>1.2), demonstrating statistically significant modulation of annotated ADRD-related pathways (**Suppl. Fig. 2b**). Gene Set Enrichment Analysis (GSEA) also showed strong upregulation of modules related to translation at synapse, RNA polymerase II regulation, and cytoskeletal organization (**Suppl. Fig. 2c**), including Tubb1/2b, Wdr34, Ssh3, Cd2ap, Nckap5l, Hap1, and nearly all members of the dihydropyrimidinase-like family (Dpysl2/3/4/5), consistent with microtubule reorganization and actin remodeling early in tauopathy (**Suppl. Fig. 2a&b**). In addition, proteins associated with mitochondrial function, including respiratory chain assembly and oxidative phosphorylation, were downregulated (**Suppl. Fig. 2c**). Key mitochondrial factors such as Opa1, Letm1, Timm50, and Tomm40, and Complex I components (Ndufa4, Acad9) were reduced (**Suppl. Fig. 2d**), consistent with mitochondrial dysfunction reported in early AD models ^17^. Additional affected pathway matching known AD pathology included multiple synaptic and neurotransmission-related pathways (**Suppl. Fig. 2c**), including downregulation of dozens of synaptic proteins involved in vesicle cycling (Syt2/7/12, Rab27b, Atg9a; **Suppl. Fig. 2e**) and neurotransmission (Gabrb2/a1, Grm8/2, Gria3; **Suppl. Fig 2f&g**).

These pattern of changes were even more pronounced at 6 months, with 253 proteins significantly upregulated and 249 downregulated (**Suppl. Fig. 3a**; **Supplementary Table 1**). GSEA revealed sustained upregulation of cytoskeleton organization, MAPK signaling, and protein synthesis, with persistent downregulation of oxidative phosphorylation, synaptic transmission, and mitochondrial calcium handling (**Suppl. Fig. 3b**).

By 9 months of age, PS19 mice exhibited widespread proteomic remodeling, but extended to 361 significantly upregulated and 353 downregulated proteins (p <0.05, FC ≥1.2; **Suppl. Fig. 3c&d, Supplementary Table 1**). In line with human AD pathology ^6,18^, they showed strong upregulation of inflammatory and astrocytic response pathways **(Suppl. Fig. 3e)**, including Apoe, Clu, C1qa, CD14, CD44, Itgb2, Serpina3n, Lyn, and HCK, and complement components (C1qa/b/c, C1ql3, C3, C4b) known to mediate synaptic pruning and promote MAPT pathology (**Suppl. Fig. 3f&g**). Conversely, we observed broad disruption of RNA metabolism and protein homeostasis, including mRNA splicing factors (Slu7), cytoskeletal components (Pfn2, Dctn6), and regulators of ubiquitin signaling (Ube2v1, Zfp91) (**Suppl. Fig. 3h; Supplementary Table 1**). Together this data reveals strong and consistent proteome changes in the PS19 mouse brain, recapitulating human AD signatures.

### Phosphoproteomics reveals MAPT-associated signaling perturbations across the disease course

Next, we performed global phosphoproteomic profiling on hippocampus and cortex samples from the same PS19 and WT mouse cohorts (**Fig. 2**). Phosphopeptides were enriched using metal ion affinity chromatography following offline fractionation, enabling sensitive and deep coverage of low-abundance phosphosites across disease progression. We identified 11,570 high-confidence phosphosites mapping onto 3,942 proteins (**Supplementary Table 2**). These included 35 unique to the human MAPT transgene and 19 to endogenous mouse MAPT. Among these, 22 phosphosites showed significant upregulation (p <0.05) beginning at 3 months that persisted through 9 months pathology (**Suppl. Fig. 1e**). These correspond to multiple phosphorylation events that are well-established for Alzheimer’s (e.g. T181, S202, S396, S404), frontotemporal dementia (e.g. S199, S202), and PSP (e.g. S202, S404), many of which are targets of stress-associated kinases such as Gsk3β, Cdk5 and Fyn. We also observed concomitant changes in phosphorylation for proteins linked to cytoskeletal disruption (**Fig. 2a-c**), synaptic and mitochondrial dysfunction (**Fig. 4d – h & Supplementary Table 2**), neuronal process collapse in early and mid-stages, and glial activation and neuroinflammatory response in the late stage (**Supplementary Table 2**). These data highlight phosphorylation events during tauopathy-associated disease course.

**Figure 2.**
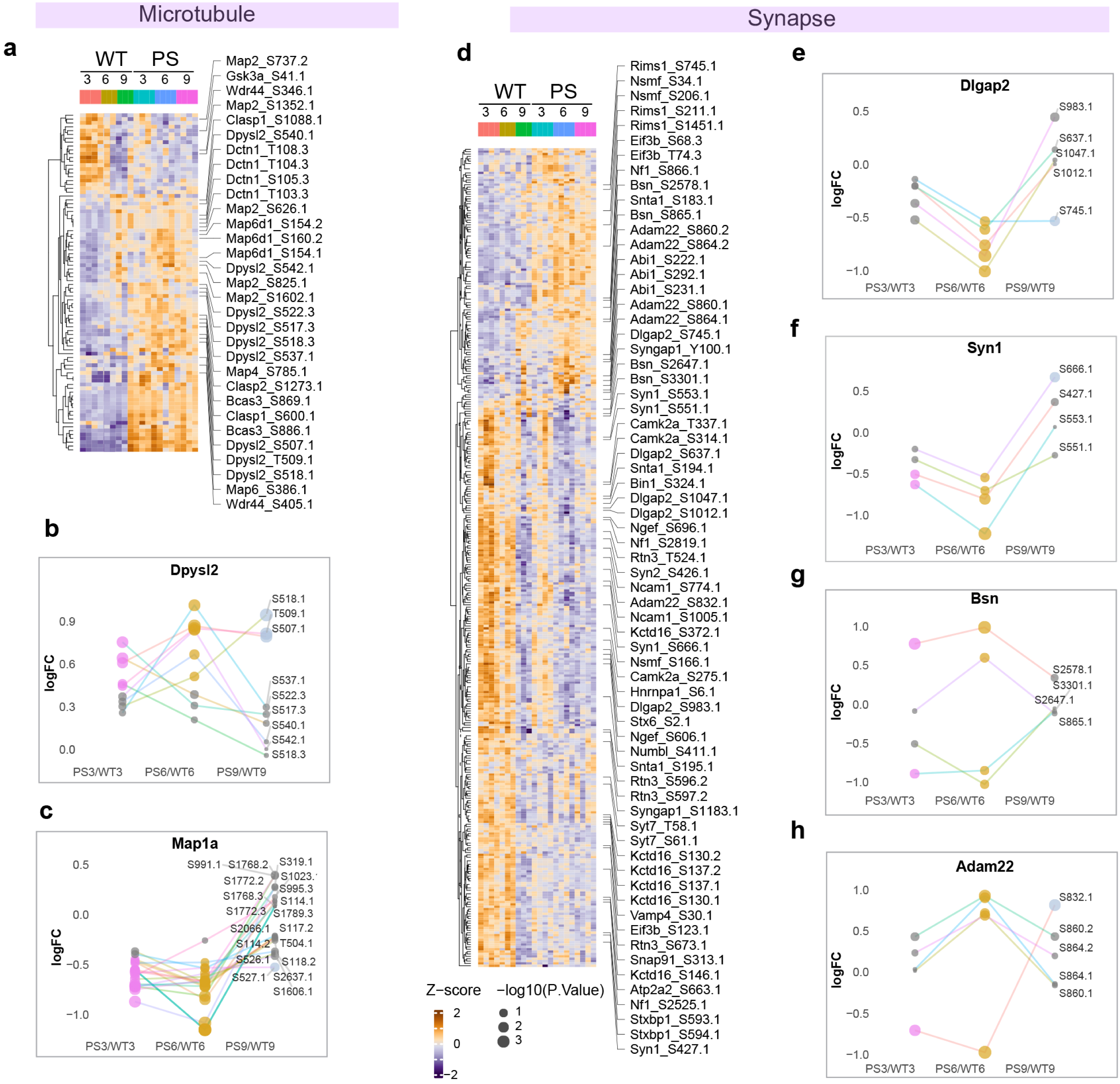
Phosphoproteomic dynamics across tauopathy progression. (**a**) Phosphorylation of MAPT and multiple cytoskeletal proteins was detected at all timepoints. (**b**) Dpysl2 phosphorylation increased at 3 and 6 months, then declined by 9 months. (**c**) Map1a phosphorylation increased at 9 months. (**d–h**) Representative phosphorylation of synaptic proteins over time, including postsynaptic markers Dlgap2/3 (**e**), vesicle-associated protein, Syn1 (**f**), presynaptic cytomatrix protein, Bsn (**g**) and the disintegrin, Adam22 (**h**).

Across the 3- and 6-month timepoints, we detected significant changes in total phosphorylation levels across the proteome of PS19 mice compared to WT, with hyperphosphorylation seen at 306 and 403 sites, respectively (PS19 *vs*. WT mice, FC >1.2, p <0.05; **Supplementary Table 2**), and reduced phosphorylation at 296 and 376 sites (PS19 *vs*. WT mice), respectively. By 9 months, the number of significantly differential phosphosites declined (192 hyperphosphorylated, 275 dephosphorylated, PS19 *vs*. WT mice; **Supplementary Table 2**). Reductions in phosphorylated proteins could have multiple causes, including synaptic and neuronal loss with disease, insolubility of aggregated proteins including MAPT (which is hyperphosphorylated) and insolubility of misfolded RNA-binding proteins. In contrast, late-stage samples showed enrichment of phosphosite changes in proteins involved in astrocytosis, microglial activation, and immune signaling. These included phosphorylation of Gfap regulators (Gap43^S142^), Bin1, Shank3, and VPS13 family proteins (Vps13a/c), which mediate lipid trafficking in glia (**Supplementary Table 2**). Dephosphorylation of Git1, Abi1/2, Jun, and Rtn4, is implicated in dendritic pruning and glial plasticity, further pointing to tauopathy-induced neuroinflammation (**Fig. 2d**). We also noted dephosphorylation of Bsn (**Fig. 2g**), which is implicated in propagation of MAPT at the synapse ^19^.

Collectively, these data reveal a temporal cascade of phosphorylation changes during tauopathy progression and highlight functionally modulated proteins in response to increasing MAPT pathology. Striking changes in phosphorylation of proteins involved in cytoskeletal maintenance and neuronal collapse, such as Mapl1a and Dpysl2 (**Fig. 2b&c**), are observed by mid-stage. Changes in hyperphosphorylation of synaptic regulators such as Dlgap2, Syn1 and Bsn are also apparent by mid-stage (**Fig. 2e, f&g**). This highlights the profound remodeling that begins early in tauopathy, presumably driven by pathology associated with MAPT-driven structural and functional decline. The late-stage shifts expand to include responses to neurodegenerative processes, these responses include inflammation, glial remodeling, and, of course, neuronal loss. These dynamic changes overlap with known MAPT kinase networks, are consistent with our prior analysis of complexes in the normal mouse brain defined by the BraInMap data set ^8,20-22^.

### Interactome profiling reveals stage-specific remodeling of protein complexes in tauopathy

As a key next step, we examined the dynamics of native multi-protein assemblies over the course of tauopathy progression by applying our optimized multiplex co-fractionation mass spectrometry (mCF/MS) pipeline ^23^ to the same PS19 mouse brain regions at 3, 6, and 9 months, again spanning early, mid, and late disease stages. Hippocampal and cortical lysates were prepared under gentle conditions to preserve multiprotein assemblies and fractionated using tandem anion and cation exchange chromatography ^8^. This setup efficiently captured both stable and dynamic complexes while minimizing denaturation. Fractions were desalted, digested, and labeled with multiplex barcoding (TMT) reagents, then analyzed by quantitative LC-MS (**Fig. 1b**).

As described for BraInMap ^8^, mCF/MS identifies numerous potential complexes. Interacting protein pairs were scored based on similarity in their chromatographic co-elution profiles. Subsequent reconstruction of multi-protein complex memberships was done using high-scoring PPIs and reproducibility across biological replicates using EPIC, a supervised machine learning tool trained on assemblies annotated in CORUM, IntAct, and GO^23,24^.

Across all timepoints and brain regions, we resolved 4072 high-confidence multiprotein assemblies (**Supplementary Table 3;** total number of complexes based on the sum across 3, 6 and 9 months, with each complex identified by a CID number followed by the age and genotype (MAPT or WT). Notably, more than half of these complexes exhibited significant changes in either abundance (p <0.05, FC >1.2, after Benjamini-Hochberg correction), composition, or connectivity during disease progression (**Fig. 3a, Supplementary Table 3),** indicating that the disease process drives broad interactome changes. Assemblies affected include those containing MAPT and its direct interactors and pathologically associated assemblies.

**Figure 3.**
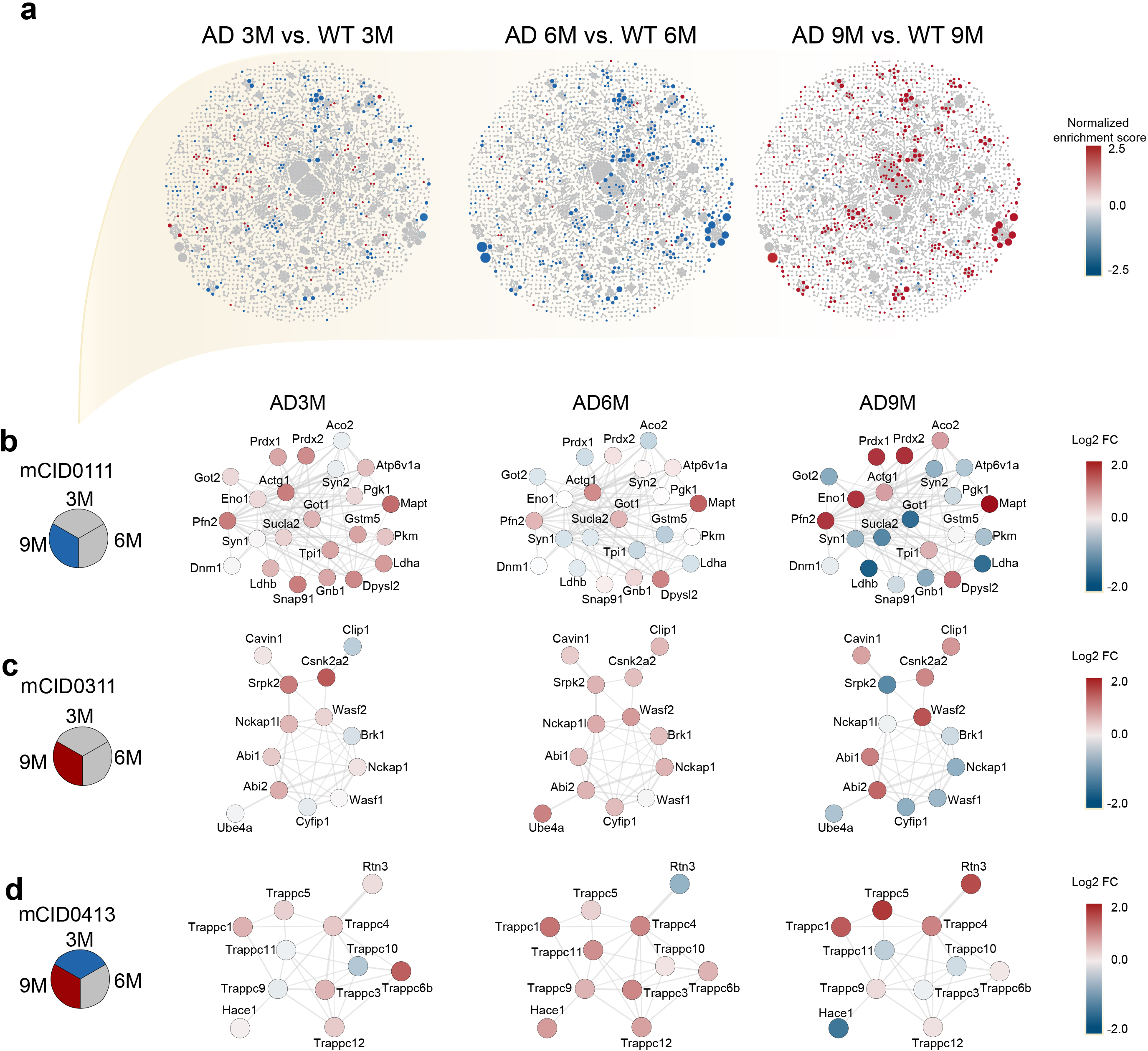
Tauopathy-associated protein complexes and interactome remodeling. (**a**) Co-fractionation and machine learning resolved 647 high-confidence protein complexes across disease stages. (**b–d**) Three representative complexes showing dynamic remodeling in tauopathy are depicted. Complex abundance changes were measured by log fold-change (PS19 vs. WT), with normalized enrichment scores (NES) used to summarize global up/down-regulation per complex based on CF/MS.

### Multi-omic data integration

Macromolecular complexes often share a common core and differ only in a subset of additional subunits. To simplify the comparison of different protein assemblies, we created a list of merged complexes that contain the same core elements, facilitating the evaluation of protein complex evolution over time and by genotype. Merging of complexes was determined using the Simpson index ^25^, with using a similarity score of 0.8 or above (Merged core complexes are identified by an “mCID” number; **Supplementary Table 4&5**). Each row in **Supplementary Table 5** describes the full set of proteins present in the specific mCID complex, however the amount of each protein and phosphorylation state of the protein can (and frequently do) differ depending on the age or genotype. The dynamics of the resulting 3419 mCID complexes are shown in corresponding heatmaps for all six conditions, with each heat map identified by its mCID number that is listed in **Supplementary Table 5** (Merged Heat Maps: https://doi.org/10.6084/m9.figshare.28914515).

After integration with global proteomic and phosphoproteomic datasets (https://doi.org/10.6084/m9.figshare.28914515; **Fig. 4a–g**), the resulting heatmaps illustrate multi-omic changes across disease progression, including phosphorylation (**Fig. 4**, columns 4–6), protein abundance (columns 7–9), and macromolecular complex dynamics inferred from CF/MS (columns 10–12). Statistical significance is denoted by white stars (columns 1–3), while heatmaps for protein levels (Prot), phosphorylation (Phos), and mCF/MS (Comp) use the same normalized scale, while merged complex fold changes (S-score) are independently normalized. Seven mCIDs containing MAPT with >10 components. The first set (mCID 97, 100, 107, 108, 111, 113) shared multiple interactors, including synaptic (Syn1, Syn2, Got1, Gstm5, Atp6v1a), mitochondria/glycolytic (Aco2, Gstm5, Ldhb, Prdx1/2), and cytoskeletal (Dpysl2, Actg1) factors (e.g. mCID 111, **Fig. 3b** & **4a**). For mCID 111, we observed early and persistent high levels of MAPT and Dpysl2 in PS19 compared to WT strains (**Fig. 3b & 4a**). The Dpysl family of proteins are known to be involved in axonal maintenance ^20^, suggesting a role for this complex in modulating axonal integrity. Other components co-evolved with disease progression, and a general decline with age, including acting binding protein Actg1, proton pump Atp6v1a, synaptic proteins such as Syn1 and Snap91, and pyruvate kinase Pkm (**Fig. 3b**). In contrast, a group of stress response proteins (including Aco2 and Prdx1&2) ^26-28^ instead showed strikingly increased association with Dpysl2/MAPT (**Fig. 3b**). Notably, the amount of phosphorylated MAPT associated with this complex increases with disease, which suggests disease-linked functional changes.

**Figure 4.**
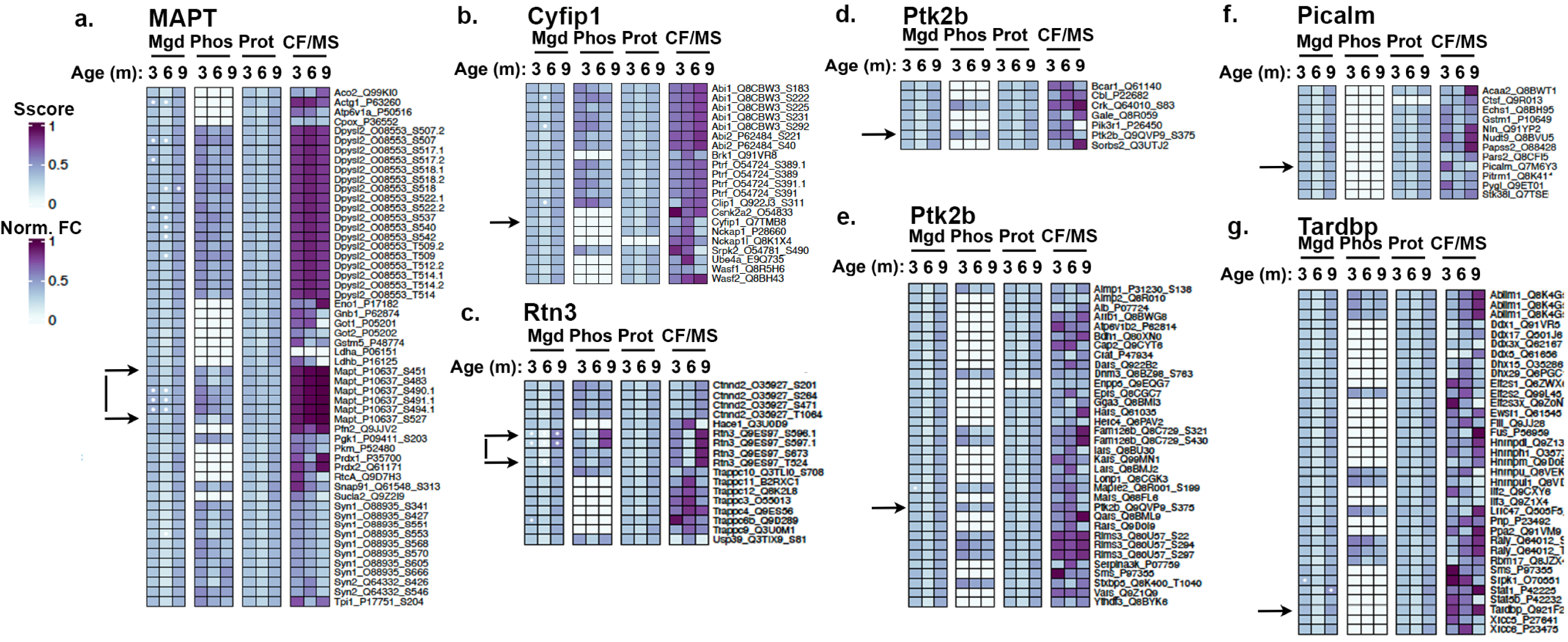
Multi-omic remodeling of tauopathy-relevant complexes. (**a–f**) Heatmaps visualize merged complex-level changes (Mgd, columns 1 – 3), across phosphoproteomic (Phos, columns 4-6), proteomic (Prot, columns 7-9) and CF/MS layers (CF/MS, columns 10-12). Key complexes include: a) Complex mCID0111 containing MAPT, b) Complex mCID0311 containing Cyfip and other actin binding proteins, c) Complex mCID0413 containing AD associated Rtn3, d) Complex mCID0884 containing AD-linked Ptk2b along with multiple known binding partners, e) Complex mCID0130 containing Ptk2b and putative binding partners suggesting a role in RNA translation and f) Complex mCID0058 containing TARDBP and multiple other RNA binding proteins. The heatmaps for Protein level (Prot), Phosphorylation (Phos) and CF/MS are normalized using the same scale, while the Merged Complex Fold Change (Sscore) was independently normalized.

Our merged complex database also reveals how other actin-binding protein (ABP) interactions change over the disease course. We focused on an assembly (mCID 311) containing several ABPs (Abi1/2, Wasf1/s), as well as the actin and FMR1 binding protein Cyfip1, which links this complex to RNA transport (**Fig. 3c** & **4b**) ^29^. mCID 311 shows a reciprocal relationship between ABPs and disease progression; the presence of Abi1 increases with disease progression whereas the presence of Abi2 drops at 6 months but rises again by 9 months (**Fig. 3c** & **4b**). Similarly, the ABP Wasf2 increases with disease progression while its homolog, Wasf1, goes down (**Fig. 3c** & **4b**), suggesting a compensatory mechanism. Additional changes implicate vesicular trafficking with disease progression as vesicular associated proteins, Clip1 and Ptrf (Cavin1), show initial increases in phosphorylation followed by increased association at higher age (**Fig. 4b**). Both factors are important for endolysosomal metabolism ^30^, suggestion its alteration leads to endolysosomal dysfunction which is increasingly understood to be an important for disease-related dysfunction and disease progression ^31^.

A third complex addresses a connection between AD pathology and vesicle trafficking at the level of endoplasmic reticulum (ER) and Golgi. mCID 413 (**Fig. 3d** and **4c**) contains multiple members of the Trappc module, which regulates ER to Golgi trafficking, indicating a vesicle trafficking function of this protein assembly. Notably, association of Rtn3 with this assembly prominently increased during disease progression (**Fig. 3d** and **Fig. 4c**). Rtn3, also localized to the ER and Golgi, is a risk factor for AD that is known to regulate the activity of BACE1 (protease that generates Aβ peptides) and accumulates in dendrites in tauopathies^32,33,34^. Our integrated analysis shows striking disease-linked increases in phosphorylation of Rtn3 at 9 months (**Fig. 2d**), not reported previously, and concomitant elevated expression (**Fig. 4c, Supplementary Table 5**). These observations point to a putative role for Rtn3 phosphorylation in ER to Golgi vesicle trafficking in AD pathology.

Multi-omics data obtained for both the ABP (mCID 311) and Trappc (mCID 413) containing complexes also highlight multi-factorial changes in the composition of macromolecules regulating vesicular/endosomal biology during disease progression, with certain subunits exhibiting alterations in phosphorylation, potentially contributing to the pathophysiology of tauopathies.

### Complexes with genetically-linked AD subunits: Ptk2b, Picalm and Tardbp

The ability to examine the composition of protein complexes across the disease course offers a powerful lens to understand how the molecular functions of disease-associated proteins evolve during progression. This is exemplified by complexes containing three Alzheimer’s–linked gene products, namely Ptk2b, Picalm and TDP-43.

The tyrosine kinase Ptk2b (Pyk2), a genetically validated AD risk factor, appears in multiple mCF-MS complexes, such as mCID 0063, 0130, and 0148, (**Fig. 4d&e; Supplementary Table 5**). Ptk2b has been implicated in MAPT pathology ^35^, Aβ production ^36,37^, and microglial phagocytosis ^38^. Two main classes of Ptk2b-containing complexes emerge. The first (e.g. mCID0884, **Fig. 4d**) contains *Crk, Cbl* and *Sorbs2*, all previously linked to Ptk2b signaling, involved in regulation of actin cytoskeletal organization and increase their association with Ptk2b with disease progression ^39-41^. Likewise, the enzyme Gale (with known mutations related to deafness and early onset cataracts shows up as a novel complex member appearing as disease progresses ^42^. In contrast, the scaffold protein *Bcar1*, another known binder of Ptk2b and Crk, is present initially but dissociates from the complex with disease (**Fig. 4d**). Decreased phosphorylation of *Ptk2b* and *Crk* (**Fig. 4d**) suggests dynamic regulation of actin cytoskeletal complexes during tauopathy.

The second class of Ptk2b-containing complexes (e.g. mCID0063, **Fig. 4e**) contains a large assembly enriched for synaptic and translational regulators, suggesting a little studied role of Ptk2b in activity-dependent synaptic translation. Ptk2b has been shown previously to regulate translation, but the mechanisms remain unknown ^43,44^. Examining this Ptk2b complex provides insight into the mechanisms through which Ptk2b might regulate translation since it includes *Aimp1/2* and a suite of tRNA synthetases (*Dars, Hars, Iars, Kars, Lars, Mars, Qars, Rars, Vars*), along with *Ythdf3*, an m6A reader that directs mRNA to synapses ^45-47^. Synaptic localization of this assembly is supported by the presence of *Rims3* and *Stxbp5*, while signaling components like *Arrb1*, *Cap2*, and *Fam126b* indicate receptor-linked pathways. Several tRNA synthetases change with disease: *Qars* and *Hars* are upregulated at 9 months, while others trend downward. These shifts suggest codon-biased translation ^48^. Phosphoproteomic data show decreased phosphorylation of *Ptk2b*, *Aimp1*, and synaptic/cytoskeletal proteins (*Mapre2, Dnm3, Rims3*) (**Fig. 4e**), supporting a phosphorylation-regulated role for this complex in synaptic remodeling.

Polymorphisms in *Picalm* are established AD risk factors, and while its gene product is implicated in vesicular and endolysosomal trafficking ^49^. Analysis of Picalm-containing complexes, such as mCID0372 (**Fig. 4f, Supplementary Table 5**), and larger assemblies including mCID0145 and mCID0434, reveals enrichment for proteins involved in energy metabolism and mitochondrial membrane biology. For example, mCID0372 includes mitochondrial enzymes such as *Acaa2* (acetyl-CoA acyltransferase 2), *Echs1* (enoyl-CoA hydratase), *Gstm1* (glutathione S-transferase), *Pitrm1* (a mitochondrial peptidase) and *Nudt9* (a NUDIX hydrolase) ^50,51^, as well as glucose metabolism enzymes like *Pygl* (glycogen phosphorylase) and *Fcsk/Fuk* (fucose kinase). It also contains tRNA synthetases (*Tars*, *Fars*, *Pars2*) with known roles in mTOR signaling and metabolism ^52,53^. Strikingly, multiple mitochondrial components become increasingly associated with Picalm as disease progresses, particularly *Echs1* and *Nudt9,* suggesting potential disease-linked roles in mitochondrial biology. This may reflect a broader function for Picalm in membrane dynamics that extends beyond vesicular trafficking and into mitochondrial maintenance during neurodegenerative stress.

Finally, we investigated TDP-43 (*TARDBP*), a key RNA-binding protein in AD, ALS, and FTD. TDP-43-containing complexes (e.g. mCID0058, 0099, 0104; **Fig. 4g**) are enriched for RBP granule proteins (*Fus*, *Ddx*, *Dhx*, *hnRNPs*, *Raly*), along with *Ablim1* and inflammatory signaling mediators *Stat1/5b* ^54,55^. Phosphorylation of *Ablim1*, *hnRNPU*, and *Raly* decreases with disease, while TDP-43 itself shows no phosphorylation, consistent with the absence of overt TDP-43 pathology in P301S MAPT mice.

### Validation by co-immunoprecipitation

To verify the composition of dynamic complexes identified by mCF/MS, we performed co-immunoprecipitation (IP) of three complex members, MAPT, Dpysl2 and Cyfip1, from brain tissue of 3-, 6-, and 9-month WT and P301S MAPT mice, followed by mass spectrometry. Although Dpysl2 has not been strongly linked to MAPT previously, both our initial and follow-up data consistently demonstrate their reproducible association. IP/MS of MAPT in 3-month P301S tissue yielded strong signal for Dpysl2, and reciprocal IP of Dpysl2 pulled down MAPT in 9-month P301S tissue. In addition, Dpysl2 IP/MS recovered shared interactors from MAPT mCF/MS complexes such as Pfkm, Nsf, Map6, and Tubb3 (**Fig. 5a, b, d**), supporting disease-relevance of Dpysl2.

**Figure 5.**
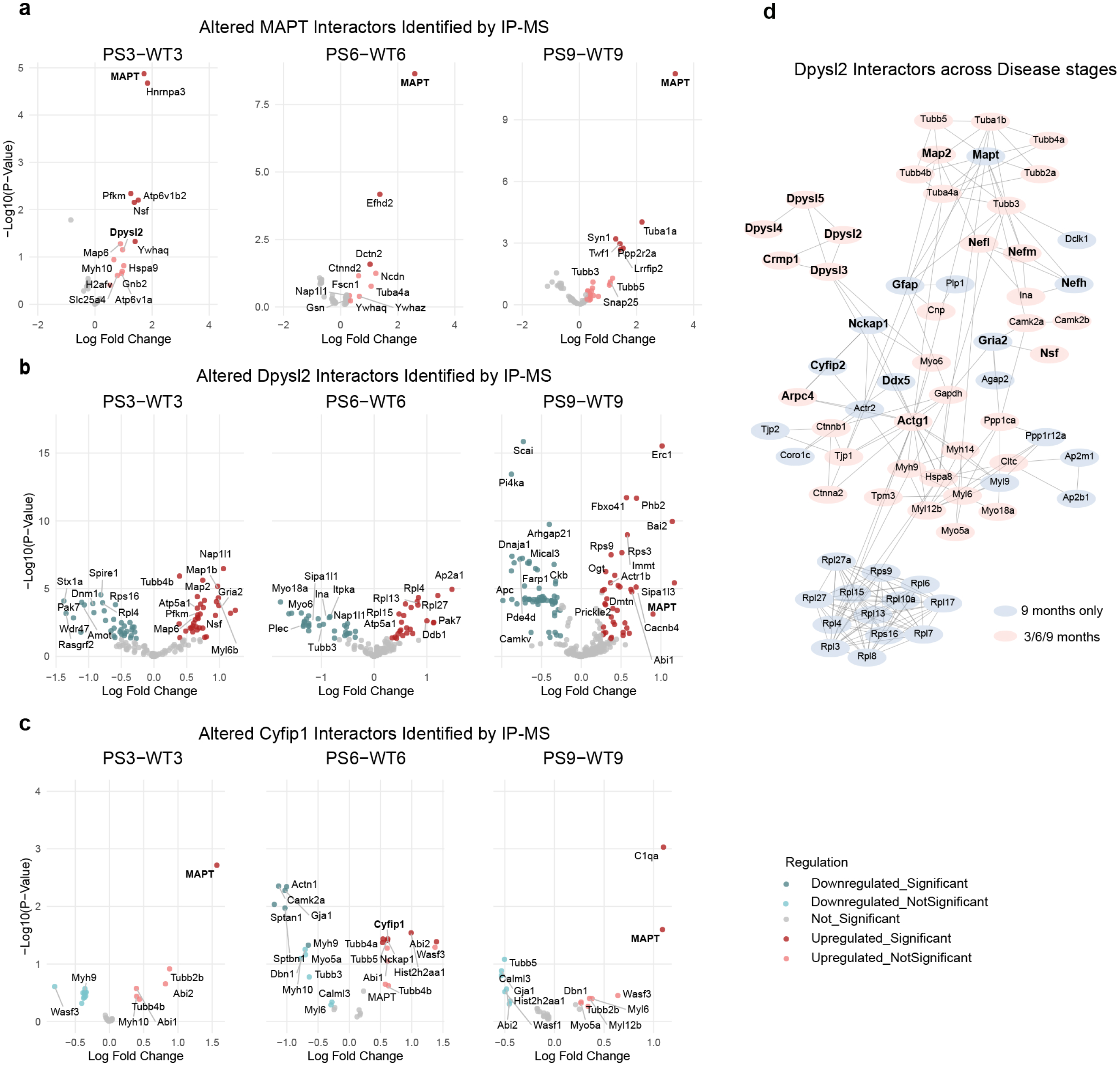
Validation of disease-linked complexes by immunoprecipitation-mass spectrometry. (**a–d**) IP-MS confirmed interactions among MAPT, Dpysl2, and Cyfip1. (**a**) MAPT pull-down recovered Dpysl2 and other CF/MS-identified interactors (e.g., Atp6v1b2, Syn1, Tubb2, Dctn2, Pfkm). (**b–c**) Reciprocal IP of Dpysl2 and Cyfip1 recovered MAPT, confirming robust interaction. (**d**) Dpysl proteins (Dpysl1–5) co-immunoprecipitated with actin regulators (e.g. Nckap1, Cyfip2, Arpc4, Actg1) and synaptic proteins (e.g. Nefl, Nefm, Nefh, Gria2, Nsf), with some interactions becoming prominent at 9 months.

MAPT IP recovered multiple previously reported MAPT interactors such as tubulins (Tuba1a, Tuba4a, Tubb3, Tubb5), Hsps, actin-binding proteins (Dctn2, Efhd2, Fscn1), and synaptic proteins (Snap25, Syn1). Moreover, MAPT IP/MS validated additional predicted interactors from the mCF/MS dataset apart from Dpysl2, including Atp6v1a, Gsn, hnRNPA3, Hspa8, and Syn1 (**Fig. 5a**), strongly implicating these proteins in MAPT pathology.

Similarly, IP/MS of Cyfip1 confirmed its association with MAPT across all timepoints, consistent with complex mCID0311 identified by mCF/MS (**Fig. 5c**). The Cyfip1 IP also enriched for actin-regulatory proteins including Wasf1/2, Abi1/2, and Nckap1 (**Fig. 7c**), in line with its established role in actin cytoskeleton remodeling^5^ ^6^.

Together, these IP/MS results validate key mCF/MS-derived complexes and support their functional relevance in tauopathy.

### Functional validation in Drosophila

Next, we performed genetic validation in established fly AD models of neurodegeneration to further investigate the interactions and assess the *in vivo* relevance of assemblies identified by mCF/ MS. Two major complexes, mCID0111 and mCID0311 (**Fig. 5b, c**) were prioritized based on their association with MAPT and their potential roles in synaptic and cytoskeletal regulation. From these complexes, we selected 12 candidate genes for functional testing classical loss-of-function alleles in heterozygosis, RNAi-mediated knockdown or over-expression in *Drosophila* lines expressing either human wild-type MAPT (2N4R) or secreted Aβ42 (**Fig. 6a-c, Supplementary Fig. 4**). We hypothesized that perturbation of subunits acting together in a complex would result in similar defects on function, and that knockdown of protein complexes identified in a mouse P301S MAPT model would show a stronger effect on a parallel MAPT fly lines than an Aβ42 *Drosophila* model.

**Figure 6.**
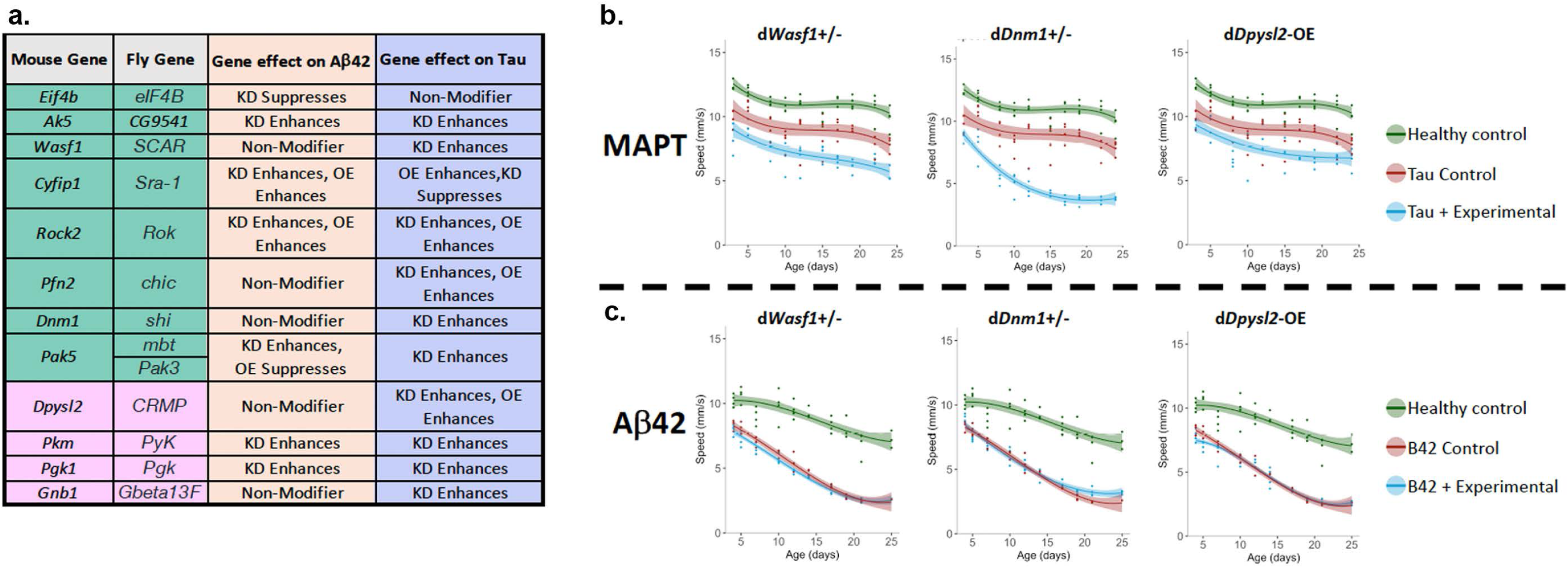
Functional validation of disease-linked complex components in Drosophila climbing assay. (**a**) Table summarizing all the functional validation experiments in Drosophila models expressing human MAPT (0N4R) or Aβ. (**b**) Climbing graphs showing the effects for the MAPT-TG crosses with Drosophila stock lines B8754 (Wasf1), B51343 (Dnm1) and B19693 (Dpysl2). (**c**) Climbing graphs showing the effects for the Aβ-TG crosses with Drosophila stock lines B10523 (Wasf 1), B30011 (Dnm1) and B19693 (Dpysl2).

The effects of candidate genes were examined in the *Drosophila* MAPT and Aβ42 models using a neuronal dysfunction assay based on quantitative behavioral (locomotor) metrics assessed longitudinally during disease progression. We found a striking difference between the results in MAPT and Aβ42 *Drosophila* models. In the MAPT-expressing flies 10 out of the 12 genes (*Ak5, Wasf1, Rock2, Pfn2, Dnm1, Pak5, Dpysl2, Pkm, Pgk1, and Gnb1*) worsened MAPT-induced deficits when knocked down; we note that 3 of these genes (*Rock2, Pfn2 and Dpysl2*) also worsened MAPT phenotypes when overexpressed. In addition, 1 gene (*Cyfip1*) worsened when overexpressed and improved MAPT-induced neuronal dysfunction when knocked down (**Fig. 6a&b, Supplementary Fig. 4**). In contrast, in the Aβ42 model, only 6 of the 12 genes (*Ak5, Cyfip1, Rock2, Pak5, Pkm,* and *Pgk1*) worsened Aβ42 phenotypes when knocked down; we note that 2 of these genes (*Cyfip1 and Rock2*) also worsened Aβ42 phenotypes when overexpressed. In addition, knockdown of *Eif4b* suppressed Aβ42-induced neuronal dysfunction (**Fig. 6a, c, Supplementary Fig. 4**).

Notably, all 11/12 complex members (92%) tested from mCID0111 and mCID0311 significantly modified MAPT-associated phenotypes, while 7 of the 12 complex members (58%) altered Aβ42-induced neuronal dysfunction, demonstrating a high degree of concordance between biochemical complex membership and functional impact *in vivo*. These independent validation experiments provide strong evidence for the disease relevance of these complexes in tauopathy and highlight their selectivity in modifying MAPT-, but not Aβ42-, driven pathology.

Together, these results support the application of interactomics to define functionally important disease modifiers and nominate modules as candidate therapeutic targets for MAPT-driven neurodegeneration.

## DISCUSSION

Our integrative survey reveals a temporally coordinated cascade of network remodeling during tauopathy, offering a high-resolution view of molecular pathogenesis not accessible from late-stage human postmortem tissue. By applying a multimodal, time-resolved approach in the PS19 tauopathy mouse model, we mapped coordinated changes in protein abundance, phosphorylation, and macromolecular complex architecture that both precede and accompany neurodegeneration. To encourage community access to this resource, the data are publicly accessible via a dedicated web portal (https://doi.org/10.6084/m9.figshare.28914515).

Focusing on specific disease-linked proteins and complexes revealed novel functional insights relevant to AD/RD. We uncovered robust and previously uncharacterized interactions between MAPT and the cytoskeletal proteins Dpysl2 and Cyfip1, which we validated by co-IP and functional assays in a *Drosophila* model system, where knockdown significantly modified MAPT-driven phenotypes. We also mapped dynamic interactomes of genetically linked AD/ALS proteins, Ptk2b, Picalm, Rtn3, and TDP-43, revealing disease-stage–specific associations with mitochondrial, synaptic, and translational machinery. For example, Picalm complexes increasingly incorporated mitochondrial enzymes as disease progressed, while Ptk2b became enriched in complexes containing synaptic tRNA synthetases and m6A regulators, pointing to a role in activity-dependent translation.

Our findings align well with previous proteomic studies of human AD/ADRD patient brain specimens ^3,6^, with late-stage enrichment of inflammation, synaptic pruning, and astrocytosis observed. However, our BraInMap data uniquely reveals extensive remodeling of the cytoskeleton and translation-related assemblies at early and mid-stages. Notably, actin-regulatory proteins, such as Actg1, Abi1–3, Dnm1, Nckap1, and Wasf1, were observed within the earliest restructured complexes, highlighting the large-scale reorganization of cytoskeletal machinery as tauopathy progresses.

The RNA translational machinery also emerged as a key early target. Complexes involving RBPs such as TDP-43 (e.g. mCID058), as well as G3BP1/2, FUS, DDX3X/5, and hnRNPs (e.g. mCID007, 013, 014) showed disease-linked compositional and phospho-remodeling. Decreased phosphorylation on key components like Ablim1, Hnrnpu, and Raly coincided with disease progression, suggesting altered phase dynamics of RNA granules. The absence of TDP-43 phosphorylation in these complexes may reflect the detection of a more soluble, dynamic pool in P301S MAPT mice, consistent with the lack of overt TDP-43 pathology. This fits with prior evidence that aggregation-prone RBPs, enriched in low-complexity domains, become sequestered in persistent stress granules during disease and are targeted for lysosomal degradation, a process impaired in aging and neurodegeneration ^56-58^.

While we observed many canonical changes in tauopathy, including increased phosphorylation of MAPT, induction of inflammatory markers such as complement C1q and Clu, loss of mitochondrial electron transport chain components, and upregulation of astrocytic markers like Gfap and S100, our co-fractionation approach revealed many additional dynamic regulatory complexity. Indeed, despite generally stable individual subunit levels, many complexes underwent dramatic shifts in membership. For instance, MAPT progressively dissociated from large synaptic assemblies while forming smaller, potentially aggregation-prone complexes. These transitions reflect both structural and regulatory changes that are largely invisible to conventional proteomics.

Integration of our interactome data with phosphoproteomics further revealed complex remodeling driven by phosphorylation of a subset of key regulatory nodes. For example, reduced phosphorylation of Syn1, Dpysl2, and Syntaxin family members implicates early disruption of vesicle trafficking and neurotransmission. These phosphosites are conserved and predicted to influence protein–protein interactions, highlighting their potential as therapeutic entry points.

Among the most striking findings was the previously unreported association between MAPT and the CRMP family (Dpysl2–5) which is involved in regulating microtubule assembly. The functional relevance of this interaction is supported by co-IP validation and by genetic interaction in *Drosophila*. Both MAPT and Dpysl2 undergo disease-linked phospho-modulation, and loss of Dpysl2 exacerbated MAPT-induced phenotypes suggests a tightly coordinated role in axonal collapse during neurodegeneration.

Additional findings from Ptk2b and Rtn3 complexes underscored the utility of mCF/MS in resolving disease-stage–specific functions. Rtn3, an AD risk gene and member of the TRAPPC trafficking complex, became phosphorylated and incorporated into complexes at 9 months in PS19 mice. Ptk2b was found in synaptic complexes enriched for translation machinery, including tRNA synthetases and the m6A-binding protein Ythdf3, implicating phospho-regulated control of synaptic translation. These results complement growing evidence that MAPT impacts translation through interactions with RBPs and ribosomes ^56,57^. Our recent unpublished work further links MAPT to m6A readers (e.g. HNRNPA2B1) and synaptic regulation of glutamatergic/GABAergic circuits, reinforcing the physiological relevance of these complexes.

Taken together, these findings establish a novel multi-omic framework for capturing the composition, dynamics, and regulatory state of protein complexes in tauopathy. Beyond identifying disease-associated proteins, our study reveals how their biochemical context shifts over time, uncovering new mechanistic insights into proteostasis, trafficking, and translational control. The observation that specific post-translational changes occur in a limited set of complex members despite widespread phosphorylation potential suggests a highly selective regulatory logic. Understanding how these molecular assemblies evolve with disease progression can inform the development of more precise therapeutic interventions.

### Limitations of the Study

Using the BraInMap workflow to study endogenous protein complexes based on biochemical co-purification necessitates a focus on soluble assemblies and mild lysis conditions. Such a focus necessarily misses at least three major types of complexes. Pathological complexes composed of misfolded, aggregated proteins are typically insoluble and thus would not be identified by the gentle mCF/MS method. Many transmembrane proteins will be spun out and lost upon removing vesicles. Finally, many nuclear proteins are caught within a web of chromatin and are also not readily available for solubilization. Another limitation of this current study is the use of a single rodent model; although the PS19 model is a well-established model, it does not completely recapitulate human disease.

### Concluding Remarks

Despite limitations, our study provides a comprehensive map of dynamic soluble protein complexes in tauopathy, revealing how molecular interactions and regulatory modifications evolve throughout disease progression. By integrating interactomic, phosphoproteomic, and functional data, we establish a systems-level framework for understanding the rewired biochemical architecture underlying neurodegeneration. These findings nominate previously unrecognized macromolecular assemblies and regulatory nodes as candidate therapeutic targets while offering a broadly applicable resource for the AD/RD research community.

## Acknowledgements

AE and BW acknowledge generous grant support from the NIH (R01AG064932, RF1AG061706) for this work. The authors thank Carl White for mass spectrometry data management.

## Author Contributions

WL, AE, and BW jointly conceived the project. WL designed and performed multi-omic experiments and data analysis; SP performed CF-MS data analysis; SJFVdS & ARO contributed to immunoprecipitation/mass spectrometry validation; NL contributed to sample preparation; RH and PH contributed to instrumentation; MCS designed and performed Drosophila studies; AKT, RR, and LJ generated and harvested mice; BJ designed Drosophila studies; WL, AE, and BW co-wrote the manuscript.

## Data and code availability

Phosphoproteomic, proteomic and mCF-MS raw data are deposited in MassIVE under accession number MSV000097878 and are publicly available. This paper does not report original code.

## Supplementary Figure Legends

**Supp. Fig. 1:**
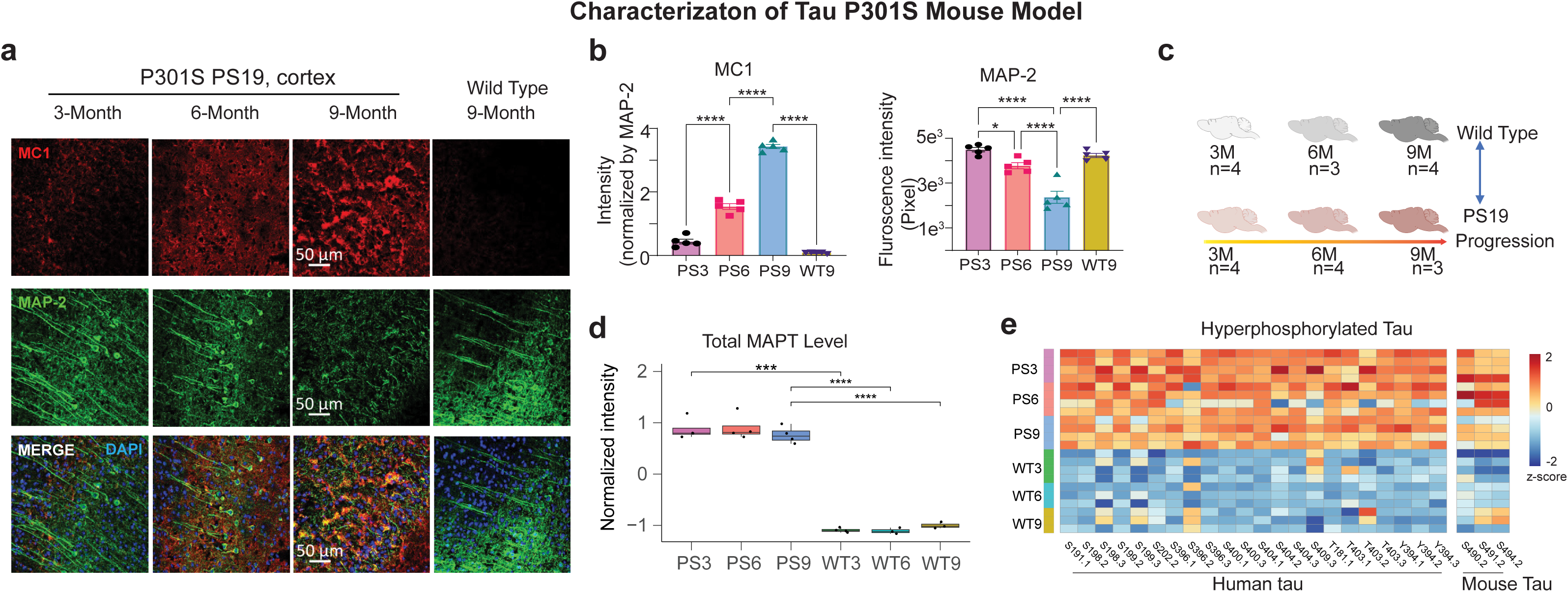
Characterization of PS19 tauopathy model and early-stage proteomic changes. (**a**) Representative immunohistochemistry of cortical sections from PS19 and WT mice at 3, 6, and 9 months, stained for misfolded MAPT (MC1, *red*) and dendritic marker MAP2 (*green*). MC1-positive aggregates were evident by 6 months and increased at 9 months in PS19 mice; no signal was observed in WT controls. (**b**) Quantification of MC1 and MAP2 staining confirmed progressive MAPT accumulation (n=5, *p<0.05).

**Suppl. Fig. 2.**
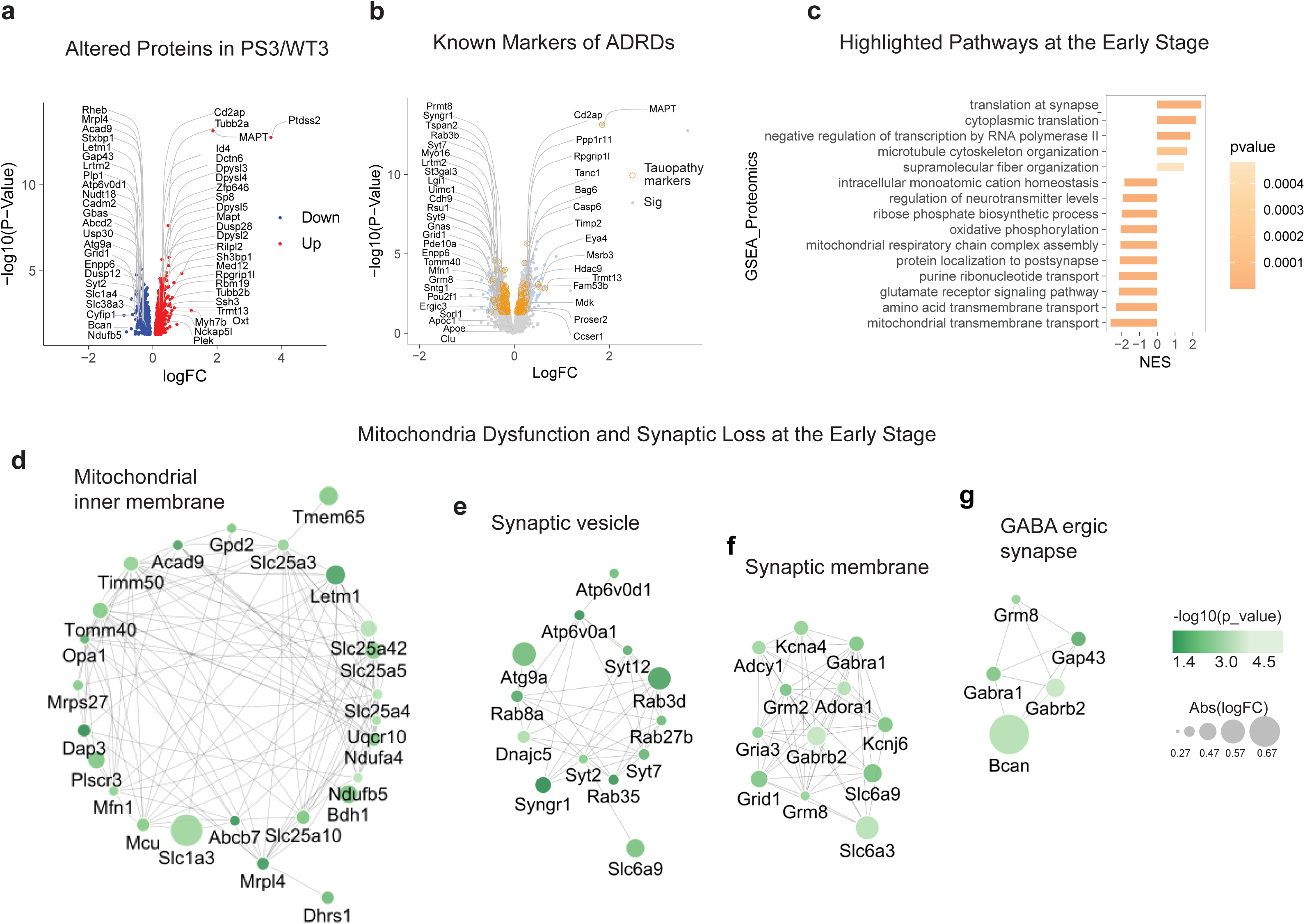
Proteome for PS19 vs. WT mice at 3 Months: (**a**) Global proteomics revealed increasing human MAPT expression in PS19 mice. (**b**) Early-stage differential expression identified key tauopathy markers. (**c**) Comparison to proteins to known ADRD-linked proteins reported by GWAS. (**d**) Enrichment analysis of early-stage changes revealed upregulation of cytoskeletal pathways and downregulation of synaptic and mitochondrial processes. (**e - h**) These included reduced expression of both mitochondrial (e.g. Tomm40, Timm50, Acad9, Opa1, Mfn1, Ndufb5) and synaptic factors (e.g. Syt2, Gria3, Grm8).

**Suppl. Fig. 3.**
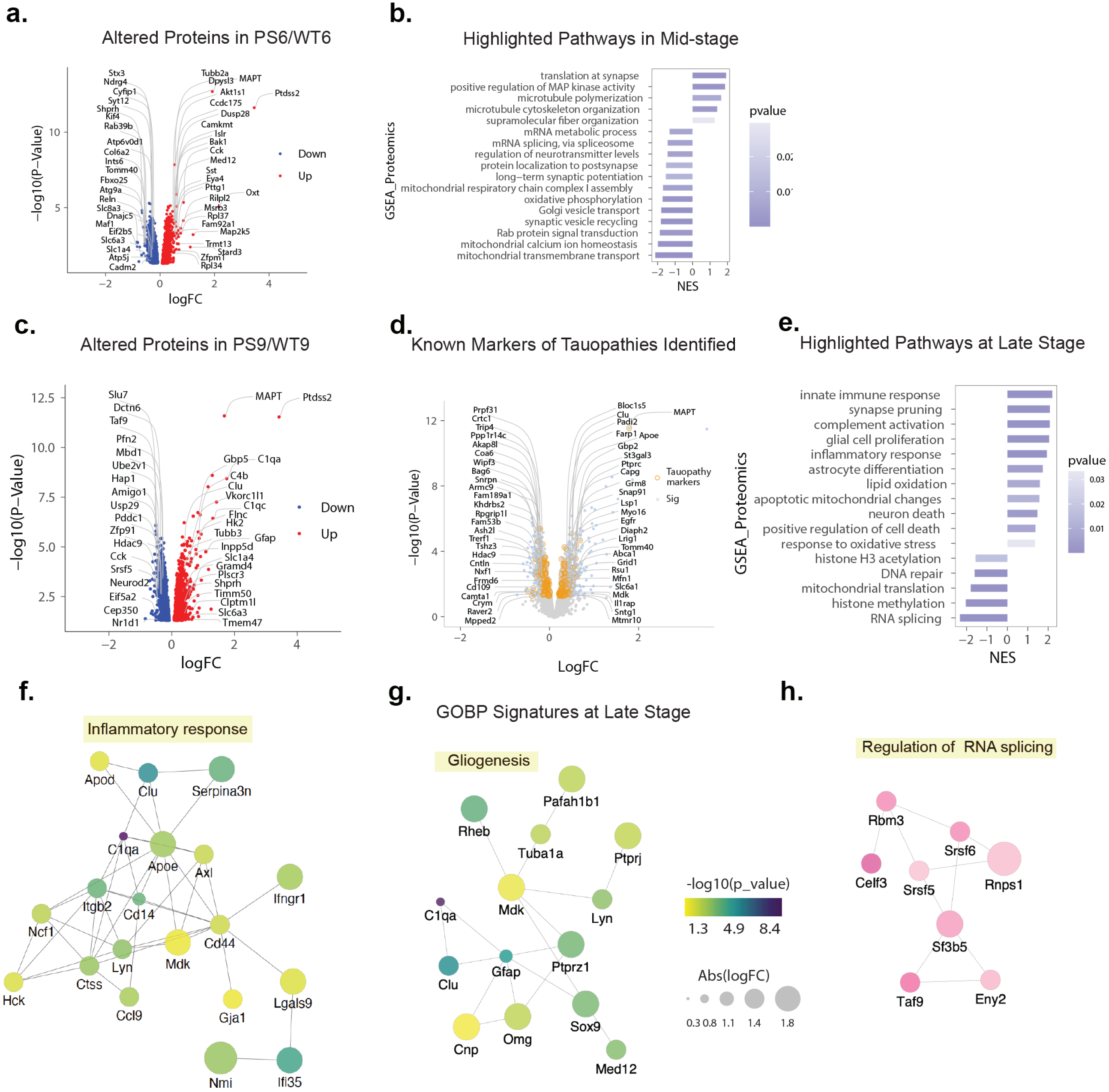
PS19 at 6 & 9 months showing proteomic remodeling at mid- and late stages of tauopathy: **(a)** Many proteins differentially expressed at 6 months overlapped with 3-month data. (**b**) Enrichment analysis of mid-stage changes (6m). (**c**) Proteomics of PS19 and WT mice at 9 months, showing upregulation of inflammatory proteins (e.g. C1qa, Clu, C4b, Tubb3, Slc6a3) and downregulation of splicing and cytoskeletal proteins (e.g. Slu7, Dctn6, Pfn2, Hdac9, Neurod2). (**d**) Comparison to proteins to known ADRD-linked proteins reported by GWAS. (**e**) Enrichment analysis for 9-month mice showing inflammatory and glial activation pathways and suppression of RNA processing. (**f-h**) Interaction mapping for inflammatory (**f**), glial (**g**) and RNA binding proteins (**h**) changed at 9 months.

**Suppl. Fig. 4.**
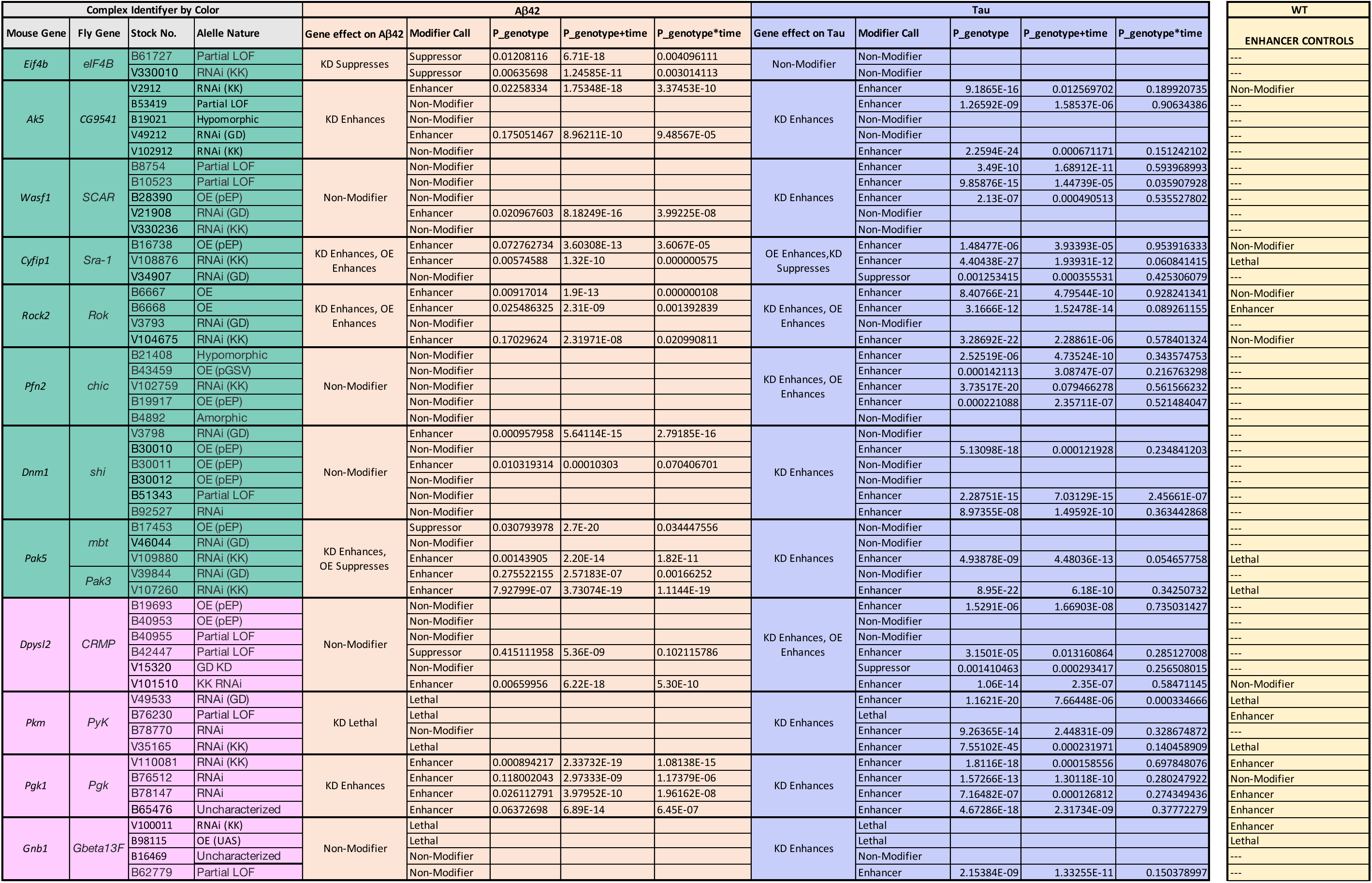
Table summarizing all the functional validation experiments in Drosophila models expressing human MAPT (0N4R) or Aβ, showing results for each Drosophila cross examined, with the statistical information for the genotype, genotype + time and genotype x time.

## STAR Methods

### Resource availability

All obtained raw data have been deposited in the MassIVE database (https://massive.ucsd.edu/) under the accession MSV000097878.

The Merged Complexes are available at the following URL: https://doi.org/10.6084/m9.figshare.28914515

### Mouse brains

The tissues used in this study for LC-MS analysis were hippocampus and cerebral cortical regions from brains of age-matched mice including 3, 6 and 9 months with wild type (n=4, male) and PS19 P301S MAPT mouse model (n = 4, male). The frozen tissues (PBS perfused) were obtained from Wolozin Lab (Boston University School of Medicine). The mice were characterized by Tau protein (MC1) and microtubule-associated protein 2 (Map2, see Supplementary Figure 1. a, b, c).

### Method details

#### Protein complexes extraction

Frozen hippocampus and cortex tissues were thawed on ice, and white matter was carefully removed. The remaining tissue was rinsed twice with ice-cold 1× PBS (ThermoFisher), then homogenized using a Dounce tissue grinder (ThermoFisher) in lysis buffer composed of 10 mM HEPES (pH 8.0), 150 mM NaCl, 5% glycerol, 0.5% n-dodecyl-β-D-maltoside (DDM), and 1 mM dithiothreitol (DTT), supplemented with protease inhibitor cocktail (Sigma) and PhosSTOP phosphatase inhibitors (Roche).

To reduce DNA contamination, 2 µL of Benzonase (Sigma) was added to the lysate, followed by incubation at 4 °C for 30 minutes with gentle rotation. Samples were then centrifuged at 12,000 × g for 10 min at 4 °C, and the resulting supernatant was collected into a fresh tube. Protein concentration was determined using the BCA protein assay kit (ThermoFisher). From each sample, 2 mg of protein complex extract was injected into an ion exchange chromatography (IEX) column for co-fractionation. Additionally, 200 µg of protein was aliquoted for proteomic and phosphoproteomic analysis, and 500 µg was reserved for immunoprecipitation-mass spectrometry (IP-MS) experiments. Remaining protein extracts were snap-frozen in liquid nitrogen and stored at −80 °C until further analysis.

#### Ion exchange fractionation of native protein complexes

Native protein complex extracts (2 mg per sample) were fractionated using an Agilent 1260 Infinity HPLC system (Agilent Technologies) equipped with tandem ion exchange columns, PolyCAT and PolyWAX (200 × 4.6 mm i.d., 5 μm, 1000 Å; PolyLC Inc., USA). Columns were equilibrated with Buffer A (10 mM ammonium acetate, ThermoFisher) for 30 minutes prior to sample loading. Separation was carried out at 17 °C using a 2-hour gradient at a flow rate of 0.45 mL/min: 5–10 min to 6% Buffer B (1.5 M ammonium acetate), 10–55 min to 20% B, 55–75 min to 60% B, 75-90 min to 100% B, followed by a 15 min isocratic hold at 100% B. Fractions were collected every minute from 6 to 103 min (96 fractions total) with a fraction collector maintained at 4 °C. Fractions were dried using a SpeedVac at 45 °C, then resuspended in 75 µL of denaturation buffer containing 6 M guanidine hydrochloride (GuHCl), 100 mM Tris (pH 8.0), 40 mM chloroacetamide, 10 mM TCEP, 1 mM CaCl**_2_**, and supplemented with protease (Sigma) and phosphatase (Roche PhosSTOP) inhibitors. Samples were heated at 95 °C for 10 minutes, cooled on ice for 10 minutes, sonicated for 10 seconds, centrifuged at 1000 ×g for 10min, and the supernatant was transferred to a new plate for automated digestion and desalting using a KingFisher system following an optimized protocol. Briefly, 75 µL of a 1:1 mixture of hydrophilic and hydrophobic beads (ThermoFisher) was added to each well, followed by ethanol to a final concentration of 50%. Beads were washed three times with 300 µL of 80% ethanol, then 4 µg of trypsin was added per well for overnight digestion at 37 °C. Digested samples were sonicated for 30 seconds, centrifuged at 10×g for 10 minutes, and the supernatant was transferred to a new plate, dried in a SpeedVac at 45 °C, and resuspended in 0.1% formic acid-2% acetonitrile for LC-MS analysis.

#### Proteomic and phosphoproteomic sample preparation

Protein extracts were denatured in 250 μL of lysis buffer containing 6 M guanidine hydrochloride (GuHCl), protease inhibitors (Sigma), and phosphatase inhibitors (Roche). The samples were heated at 95°C for 10 min, cooled on ice for 10 min, and then briefly sonicated to shear the nucleic acids. The samples were diluted with 100 mM Tris (pH 8.5) to reduce the concentration of GuHCl to 0.75 M. After quantification with a BCA kit (Thermo Scientific), the proteins were digested overnight with sequence-grade trypsin (enzyme-to-protein ratio of 1:50) at 37°C, and formic acid was then added to obtain a final concentration of 1% in solution. The resulting peptides were desalted using a C18 Sep-Pak (Waters) according to the manufacturer’s instructions.

#### TMT peptide labeling

Prior to tandem mass tag (TMT) labeling, peptide quantification was performed by Pierce quantitative colorimetric assay (Thermo Scientific). According to the manufacturer’s instructions, 100 μg of peptide per sample was resuspended in 0.1 M triethylammonium bicarbonate (TEAB). Peptides of cancer cell (5 channels per condition, glucose & no glucose) were labeled with 0.2 mg of TMT 10-plex while 15 samples of human serum (5 channels per condition, healthy, pre- and post-dialysis) were labeled with TMTpro (Thermo Scientific) for 1 h at room temperature. To quench the reaction, 5% hydroxylamine was added to each sample, and the resulting mixture was incubated at room temperature for 15 min. After labeling, equal amounts of each sample were combined in a new microtube and desalted using a C18 Sep-Pak (Waters).

#### High-pH reversed-phase peptide fractionation

Peptides (500 μg) were fractionated offline on a Waters XBridge BEH C18 reversed-phase column (3.5 μm, 4.6 ×250 mm) using an Agilent 1100 HPLC system operated at a flow rate of 0.45 mL/min with two buffer lines: buffer A (consisting of 0.1% ammonium hydroxide-2% acetonitrile-water) and buffer B (consisting of 0.1% ammonium hydroxide-98% acetonitrile, pH 9). The peptides were separated by a gradient from 0% to 10% B in 5 min followed by linear increases to 30% B in 23 min, to 60% B in 7 min, and then 100% in 8 min and maintained at 100% for 5 min. This separation yielded 48 collected fractions that were subsequently combined into 12 fractions and then evaporated to dryness in a vacuum concentrator. The peptides (2 µg) from each fraction were reconstituted in 0.1% formic acid and maintained at -80°C prior to analysis by nLC-MS/MS.

#### Phosphopeptide Enrichment

Phophopeptide were enriched using Fe-NTA magnetic beads (CubeBiotech). Briefly, the Fe-NTA beads were washed three times with binding buffer of 0.1% TFA-80% ACN three times. The peptides were dried and resuspended in binding buffer then incubated with the Fe-NTA bead slurry for 30 min on a shaker. Bound phosphopeptides were washed 3 times with binding buffer, then serially eluted twice by adding 200 μL of 2.5% ammonium hydroxide-50% ACN to the beads. Eluted phosphopeptides were combined and dried in a speedvac prior to LC/MS.

#### Quantitative LC-MS analysis of peptides

LC-MS analysis was performed using the same Q-Exactive HF system as described above. Peptides were loaded onto a C18 pre-column (75 μm i.d. × 2 cm, 3 μm, 100 Å, ThermoFisher Scientific) and then separated on a reverse-phase nano-spray column (75 μm i.d. × 25 cm, 2 μm, 100 Å, ThermoFisher Scientific) using gradient elution. Peptides (2 μg) were injected and separated over a 150 min gradient. The mobile phase A was consisted of 0.1% FA-2% ACN-Water, and mobile phase B was consisted of 0.1% FA-80% ACN-Water. The gradient consisted of 6% to 40% mobile phase B over 155 min, was increased to 95% mobile phase B over 4 min, and maintained at 95% mobile phase B for 3 min at a flow rate of 250 nL/min. The MS instrument was operated in positive ion mode over a full mass scan range of m/z 350−1400 at a resolution of 60,000 with a normalized AGC target of 300%. The source ion transfer tube temperature was set at 275°C and a spray voltage was set to 2.5kv. Data was acquired on data dependent acquisition (DDA) mode and the top 10 most abundant ions were selected for fragmentation. MS2 scans were performed at 45,000 resolution with a normalized collision energy 34. Dynamic exclusion was enabled using a time window of 60 s.

#### Proteomic data analysis

MS2 spectra were processed and searched by MaxQuant (version 1.6)^59^ against a database containing native (forward) human protein sequences (UniProt) and reversed (decoy) sequences for protein identification. The search allowed for two missed trypsin cleavage sites, variable modifications of methionine oxidation, and N-terminal acetylation. Both carbamidomethylation of cysteine residues and TMT (peptide N-termini, K residues) were set as fixed modifications. Precursor ions were searched with a maximum mass tolerance of 4.5 ppm and fragment ions with a maximum mass tolerance of 20 ppm. The candidate peptide identifications were filtered assuming a 1% FDR threshold based on searching the reverse sequence database. Quantification was performed using the TMT reporter on MS2 (TMT10-plex). Proteins with less than 30% missing value across all samples were kept for quantification. The reporter ion intensities were log-transferred and normalized using quantile normalization. Bioinformatic analysis was performed in the R statistical computing environment (version 4.3.0) using a built-in tool^60^. Plots were generated using R programming language. Enrichment analysis was performed using the Gene Ontology database^61^.

#### Interactomic data analysis

The co-fractionation raw files were searched in MaxQuant v1.6.0.16^62^ against the Mus musculus protein sequence database downloaded from UniProt on October 10, 2020. Carbamidomethylation of cysteine was set as a fixed modification while variable modifications were oxidation of methionine and acetylation of protein N-termini. Two missed cleavages were allowed and reporter ion MS2 was used for quantification with 16plex TMT and a reporter mass tolerance of 0.003 Da. Peptide search tolerance for MS1 and fragment tolerance for MS2 were set to 4.5ppm and 10ppm respectively with match between runs activated at a match window of 0.7 min and an alignment window of 20 min. The false discovery was controlled using a target/decoy approach with a false discovery level set to 1%. This resulted in the detection of 4,162 proteins.

The MS1 intensities obtained through database searching were run through the EPIC prediction tool^24^ using default settings to predict PPI and complexes for each timepoint in MAPT genotype and WT samples. The results from each fractionation were processed by random forest classifier trained on experimentally verified PPIs derived from 576 complexes extracted from public curated databases to assign PPI confidence scores (CORUM, IntAct and GO). After generating a high-confidence co-elution network, we used ClusterONE^63^ to generate a set of stable protein complexes from the co-elution network. Using two-pronged approach, we predicted 39,828 PPI (**Supplementary Table XX**) leading to 4,072) complexes (**Supplementary Table 3)** across the 6 datasets (sum across 3, 6, and 9 months, with each complex identified by a CID number followed by the age and Genotype (MAPT or WT)). Since many complexes shared a common core and differed only in a subset of subunits, we further merged the complexes containing the same core elements over time and genotype if their Simpson coefficient ≥ 0.8 to get a set of 3,419 merged complexes (**Supplementary Table 4)**. To further study the differences in protein expression across complexes over timepoints we performed GSEA using proteomic protein expression profiles with merged complexes as a gene-set comparing genotype against WT for each timepoint (**Fig5a-d**).

#### Co-immunoprecipitation assay

Protein complex extracts (500 µg) from 3-, 6-, and 9-month-old wild-type or PS19 mice, obtained from the previous step, were thawed on ice and incubated overnight at 4 °C with rotation in the presence of specific antibodies. Antibodies used for Co-IP included mouse anti-tau5 (N.K., MSU), rabbit anti-Dpysl2 (Abcam, ab129082), rabbit anti-Cyfip1 (Abcam, ab172730), mouse IgG (I-2000-1, Vector Laboratories) and rabbit IgG (Abcam, ab172730). A/G agarose beads were washed twice with lysis buffer containing 0.1% n-dodecyl-β-D-maltoside (DDM), then incubated with the antibody-protein complexes for 1 hour at 4 °C with rotation. After incubation, samples were centrifuged at 1000 × g for 2 minutes at 4 °C. The beads were washed once with 0.1% DDM lysis buffer, followed by two washes with 50 mM ammonium bicarbonate. During the final wash, samples were transferred to fresh tubes, centrifuged, and the supernatant was removed. Bead-bound proteins were digested by adding 60 µL of 50 mM ammonium bicarbonate containing 1 µg trypsin and incubating overnight at 37 °C with rotation. Digestion was quenched by adding formic acid to a final concentration of 1%. Peptides were then desalted using C18 Sep-Pak cartridges (Waters) according to the manufacturer’s instructions prior to LC/MS analysis.

#### Drosophila models characterization and neuronal dysfunction assay

The UAS-Tau and UAS-β42 Drosophila lines used in this study have been previously characterized ^7^. We used the elav-GAL4 driver line to achieve neuron-specific expression of UAS transgenes. Genetic strains targeting *Drosophila* homologs of genes of interest were sourced from the Bloomington *Drosophila* Stock Center (BDSC) and the Vienna *Drosophila* Resource Center (VDRC). Homologs were identified using DIOPT (DRSC Integrative Ortholog Prediction Tool, Version 9)^8^. We included all *Drosophila* homologs with a DIOPT score ≥10 in our functional assays. For genes with scores <10, we only selected the highest-scoring homologs.

We employed a custom-built automated system to quantify neuronal dysfunction in *Drosophila* via the startle-induced negative geotaxis assay. To assess age-related motor decline, we collected 10 age-matched virgin females per genotype, with four technical replicates per genotype. Female flies were selected due to their larger size and reduced variability in motor performance during early adulthood, which enhances tracking consistency. Flies were collected on Day 1 post-eclosion and maintained on 300 μL of fresh media, with behavioral testing conducted every other day throughout their lifespan. During each session, flies were tapped to the bottom of a vial and their climbing behavior was recorded over a 7.5-second interval using our automated tracking software. Each replicate underwent three trials per testing day. Data from four replicates per genotype were analyzed using a linear mixed model to assess statistical differences between control groups (Tau or β42 alone) and experimental groups (candidate gene perturbation combined with Tau or β42)., assessing significance across three metrics; genotype, the additive effect of genotype + time, and the interactive effect of genotype ξ time.

